# Tactile Imagery Affects Cortical Responses to Vibrotactile Stimulation of the Fingertip

**DOI:** 10.1101/2023.06.02.543456

**Authors:** Marina Morozova, Lev Yakovlev, Nikolay Syrov, Gleb Perevoznyuk, Mikhail Lebedev, Alexander Kaplan

## Abstract

Although imagery of tactile sensations is not so well studied compared to other types of mental imagery, it is potentially very useful for brain computer interfaces (BCIs) where it could produce neural modulations needed for BCI operations. Here we assessed neural modulations associated with tactile imagery (TI) by comparing its effects on cortical responses to the effects of actual vibrotactile stimulation of the fingertip. We found that both TI and vibrostimulation evoked event-related frequency changes of the electroencephalographic (EEG) activity. Moreover, TI affected somatosensory evoked potentials (SEPs) evoked by short pulses of vibration. EEG data were collected in 29 participants trained to perform tactile imagery task. Responses to vibratory pulses were measured with and without TI. These SEPs consisted of three prominent components: a P100 response in the centro-parietal regions, a P200 response in the frontal region, and a P300 response in the central regions. The TI consistently resulted in an increase in ipsilateral P100, ipsi- and contralateral P300 and frontal P200. Moreover, TI strengthened the θ-band ERS in the frontal region that occurred in response to vibration. These findings suggest that TI not only modulates EEG patterns by itself but also affects cortical processing of physical somatosensory stimuli. Such conjoint processing of both real and imagined somatic sensations could be utilized in BCIs, particularly in clinically relevant BCI that strive to restore somatosensory processing by combining centrally-induced and peripheral activities.

**Significance Statement:** While it is known that tactile imagery (TI) engages the same cortical areas that are active during the processing of real tactile inputs, neural mechanisms of such shared representation are not well understood. This study employed EEG recordings to examine the interaction between real and imagined somatic sensations. It was found that TI both changes EEG oscillatory activity and facilitates cortical responses to real tactile stimuli. Therefore combining TI with tactile stimulation could be useful for tactile-based brain-computer interfaces (BCIs), particularly the ones of clinical utility for neurorehabilitation and sensory substitution.

## Introduction

Human ability to imagine events, actions and perceptions (Farah, 1989; Nanay, 2018) is both useful as a method in neurophysiology (Decety, 1996; Crammond, 1997; Jeannerod and Frak, 1999; Pylyshyn, 2002) and a practical approach in the clinic (Warner and McNeill, 1988; Van Leeuwen and Inglis, 1998; Zimmermann-Schlatter et al., 2008; Di Nuovo et al., 2014; Pearson et al., 2015). In the sensory domain, it has been repeatedly demonstrated that the same neural structures are activated by real sensory inputs and imagined perceptions (Kosslyn et al., 2001). The research on this nontrivial finding has primarily focused on the visual perceptions while other sensory modalities have not been thoroughly examined (McNorgan, 2012; Pearson, 2019). Particularly, many studies have scrutinized motor imagery (Lotze and Halsband, 2006; Zimmermann-Schlatter et al., 2008; Ladda et al., 2021) but relatively a few have examined imagined somatosensory perceptions (Yoo et al., 2003; Schmidt et al., 2014; Yao et al., 2022a; Yakovlev et al., 2023).

Our research on the electroencephalography (EEG) patterns associated with somatosensory imagery is motivated by the idea of incorporating such imagery in the design of brain-computer interfaces (BCIs) (Yao et al., 2022a; Yakovlev et al., 2023). The feasibility of this idea is supported by the results of neuroimaging studies showing that somatosensory imagery activates the same brain areas as the ones normally activated in somatosensory tasks (Yoo et al., 2003; Schmidt et al., 2014; de Borst and de Gelder, 2017; Schmidt and Blankenburg, 2019), which is similar to the activation of the matching sensory areas during imagery with other modalities. Additionally, activation of the somatosensory cortex during tactile imagery was confirmed using intracranial single-neuron recordings in humans (Bashford et al., 2021).

EEG correlates of somatosensory imagery are of particular interest because of their utility in BCI applications (Yao et al., 2022a). The event-related desynchronization/synchronization (ERD/S) of sensorimotor rhythms (SMR) in EEG (mu and β) has been widely used as a reliable marker of cortical sensorimotor processing (Pfurtscheller and Neuper, 1997; Neuper et al., 2006), including plasticity of sensorimotor system (Boonstra et al., 2007; Freyer et al., 2013). In particular, ERD reflects activation of the sensorimotor network (Klimesch et al., 2007) where low-frequency cortical oscillations are suppressed (Pineda, 2005). ERD often occurs during the execution of sensorimotor tasks in the hemisphere contralateral to the working limb (Hari and Salmelin, 1997).

Somatosensory evoked potentials (SEPs) are a useful tool for studying sequential processing of tactile information. The early components of SEPs with the latency below 150 ms are associated with the activation of the primary somatosensory cortex (S1) (Hoechstetter et al., 2001), and the longer-latency SEP components contain contributions from the secondary somatosensory areas (Karhu and Tesche, 1999). Functionally, the primary somatosensory cortex performs early tactile processing, including perception and categorization (Zainos et al., 1997). Both primary (Borich et al., 2015) and secondary (Huttunen et al., 1996) have a role in sensorimotor integration. The secondary somatosensory cortex is important for tactile attention (Mima et al., 1998), processing of the temporal features of somatic sensation (Ferrington and Rowe, 1980; Sinclair and Burton, 1991), tactile learning and memory. Additionally, S2 contributes to the initial processing of somatosensory input which it receives directly from the thalamus (Luijten et al., 2020). S2 has larger receptive fields compared to S1 which are often bilateral. Overall, the involvement of different somatosensory areas in imagery is not well understood. Barsalou (2008) and Schmidt et al. (2014) described the role of S1 in tactile imagery as perceptual grounding, and Yoo et al. (2003) reported that S2 is active during imagery, as well.

The effects of somatosensory imagery on cortical responses to physical somatosensory stimuli are of particular interest to the BCI field. This is because such measurements could be used to quantify the imagery and also because BCIs could use a combination of somatosensory stimulation and imagery. For the motor systems, similar interactions have been studied as the effects of motor imagery on the movements evoked by transcranial magnetic stimulation (TMS) (Aono et al., 2013; Takemi et al., 2013; Vasilyev et al., 2017) and spinal reflexes (Cowley et al., 2008; Takemi et al., 2015). Yet, compatible methods have not been developed for the somatosensory system. Accordingly, this study is an investigation of how tactile imagery (TI) modulates cortical responses to vibrotactile stimulation. We observed an interaction of imagined and real somatic sensations at the cortical level, which could be utilized in BCIs.

## Materials and Methods

### Subjects

Twenty nine healthy volunteers (9 females, 25.1 ± 4.8 y.o., right-handed) participated in one experimental session lasting up to 90 minutes. All participants gave written informed consent to participate in the study. The study adhered to the Human Subject Guidelines of the Declaration of Helsinki and was approved by the Ethics Committee of the Skolkovo Institute of Science and Technology.

### Experimental Design

The experimental session included 2 general parts: a learning session where subjects were trained on TI and the session where TI was combined with vibrostimulation.

During the learning part, subjects were familiarized with vibrostimulation of the right index fingertip. They were asked to mentally reproduce the sensations evoked by vibration. For this purpose, 8-s trains of randomly patterned vibration were utilized. The stimulus randomness reduced tactile habituation and helped to avoid residual tactile sensations. The training session consisted of 15 trials with vibrostimulation intermixed with 15 trials of the same duration but without the stimulation. When learning tactile imagery, participants imagined a vibrostimulation pattern for 8 s in the absence of actual stimulation. The number of TI-learning trials was 30 (arranged in 2 runs like the TS-learning trials). This approach to TI learning is described in detail in Yakovlev et al. (2023).

During the TI session with short vibratory stimuli incorporated, vibrostimulation lasting 75 ms and occuring at the intervals of 1-2 s were used. This session consisted of 2 experimental conditions: the resting state and TI arranged as the separated runs. During the resting state, participants were asked to count elements of a visual scene and pay attention to the skin surface where vibratory pulses were applied. The resting-state runs were performed in the beginning and in the end of each experimental session. For these 2 runs, the number of vibrotactile stimuli was 200 (100 per run). During the TI condition, the participants imagined being stimulated for 10 s while being cued by a vibration pictogram that appeared on the screen. 5 short vibrational stimuli were delivered at random times during the 10 s period of TI with interstimulus intervals of 1-2 s. The total number of vibrational stimuli during the TI condition was 100.

### EEG Recordings

Throughout the experiment, EEG data were collected using an NVX-52 amplifier (MKS, Russia). Forty seven EEG channels were recorded according to the international “10–10” system with the sampling frequency of 500 Hz using Ag/AgCl electrodes lubricated by an electrode gel. The ground lead was attached to the FCz site. Two reference electrodes were placed on the left and right mastoids in the positions of TP9 and TP10 ?hannels. The skin-electrode impedance was kept below 15 kΩ. During the experiment, participants were asked to keep their eyes open, sit still in a comfortable position and not to make unnecessary movements while EEG recording was in progress. The stimuli for visual attention were presented on the monitor placed at the distance of 1.2 m in front of the participant. The participants were asked to count random elements of the visual cues when they were present on the screen. Vibration was applied to the index fingertip of the right hand.

### Data Analysis

For EEG preprocessing, several channels (no more than 5) were excluded as too noisy. The data for these channels were linearly interpolated. Eye-movement artifacts were removed using Independent Component Analysis (ICA) using Fp1-Fp2 channels as EOG. The preprocessed EEG signals were band-pass filtered using a 4th order Butterworth filter in two frequency ranges: a) 0.2-12 Hz for ERP analysis in time domain, and b) 0.2-40 Hz for time-frequency analysis using the continuous Morlet wavelet transform (CWT).

For processing of records with long (8-s) vibrotactile stimulation or TI without intermittent vibrotactile stimuli, EEG data were split into 10-s epochs starting 1 s before the beginning of the segment and ending 1 s after the segment. Each epoch was z-score standardized for each channel, CWT was applied to each epoch and resulting powers of TS epochs were baseline corrected by the powers of rest epochs by subtracting the mean of rest powers followed by dividing by the mean of rest powers (i.e., percent baseline correction).

For the processing of the recordings where short (75 ms) vibrotactile pulses were applied, EEG data were split into 1.5 second long epochs starting 0.5 second before and ending 1 second after the vibration stimulus onset. Each epoch was z-score standardized for each channel. For SEP analysis, we defined approximate latency and localization of P100, P200 and P300 peaks on grand average SEP and topomaps. We calculated amplitudes of P100, P200 and P300 for each participant in each condition as average of the values in the predefined temporal-spatial range (±15 ms around grand average peak latency; chosen target leads). Resulting mean amplitudes were compared across participants using Wilcoxon signed rank test. For time-frequency analysis of the 1.5-s epochs, CWT was applied to each epoch and the resulting powers were percent baseline corrected by the average powers in the (−0.5, 0) second time period. In the resulting time-frequency maps, we defined approximate latency and localization of spectral peaks and calculated their median powers in the predefined temporal-spatial-frequency range. For pairwise comparison of the resulting mean powers, Wilcoxon signed rank test was used.

We defined the individual frequencies of μ- and β-rhythms for each participant based on the visual inspection of the spectra. The frequency of μ- and β-rhythms was then more precisely calculated as the median value in the ±1 Hz window for the 8-s and 1.5-s wavelets.

### Code/Software

Visual stimuli were delivered using PsychoPy 2022.2.5. The post hoc preprocessing and analysis was performed using Python data processing packages, including MNE 1.3.1 and SciPy 1.10.0.

We used a custom designed and computer-controlled vibrotactile stimulator based on an Arduino UNO (Arduino, Spain). A flat vibration motor (6 mm in diameter, max speed of 12,000 rpm) was placed on the index finger of the right hand. During the long-lasting (8-s) vibration, the stimulation pattern was randomly patterned: the motor speed varied in range 6,000-1,2000 rpm. Such a random stimulation pattern reduces tactile habituation and helped to avoid residual tactile sensations. During the short (75 ms) vibratory pulses, motor speed was set at 9,000 rpm.

The code for stimuli presentation and data processing is available on Github (https://github.com/MarkaMorozova/Tactile-Imagery). Full dataset generated and analyzed for this study is available upon request. Partially processed data (ERP and Time-Frequency data) are available at https://bit.ly/43tWQc3.

## Results

### Even-Related Desynchonization (ERD) during Tactile Imagery Learning

During the 8-s Tactile Stimulation (TS) trials, we observed μ- and β-rhythm ERD that were the most pronounced in the contralateral hemisphere, in the C3 and CP3 leads (Fig. 2A, Fig. 2C, Table 1). During TS, these ERDs were also noticeable in the ipsilateral hemisphere, in the C4 and CP4 leads, but the μ-rhythm ERD in the ipsilateral hemisphere was statistically significantly lower than in the contralateral hemisphere (*W* = 107, *p* = 0.0157).

**Table 1.** Powers of μ-rhythm ERD (median [25th – 75th percentiles]).

| Condition | Contralateral, % | Ipsilateral, % |
| --- | --- | --- |
| Tactile Stimulation | -26 [-51 – -14] | -19 [-34 – -11] |
| Tactile Imagery | -21 [-41 – -3] | -23 [-39 – -2] |

**Figure 1.**
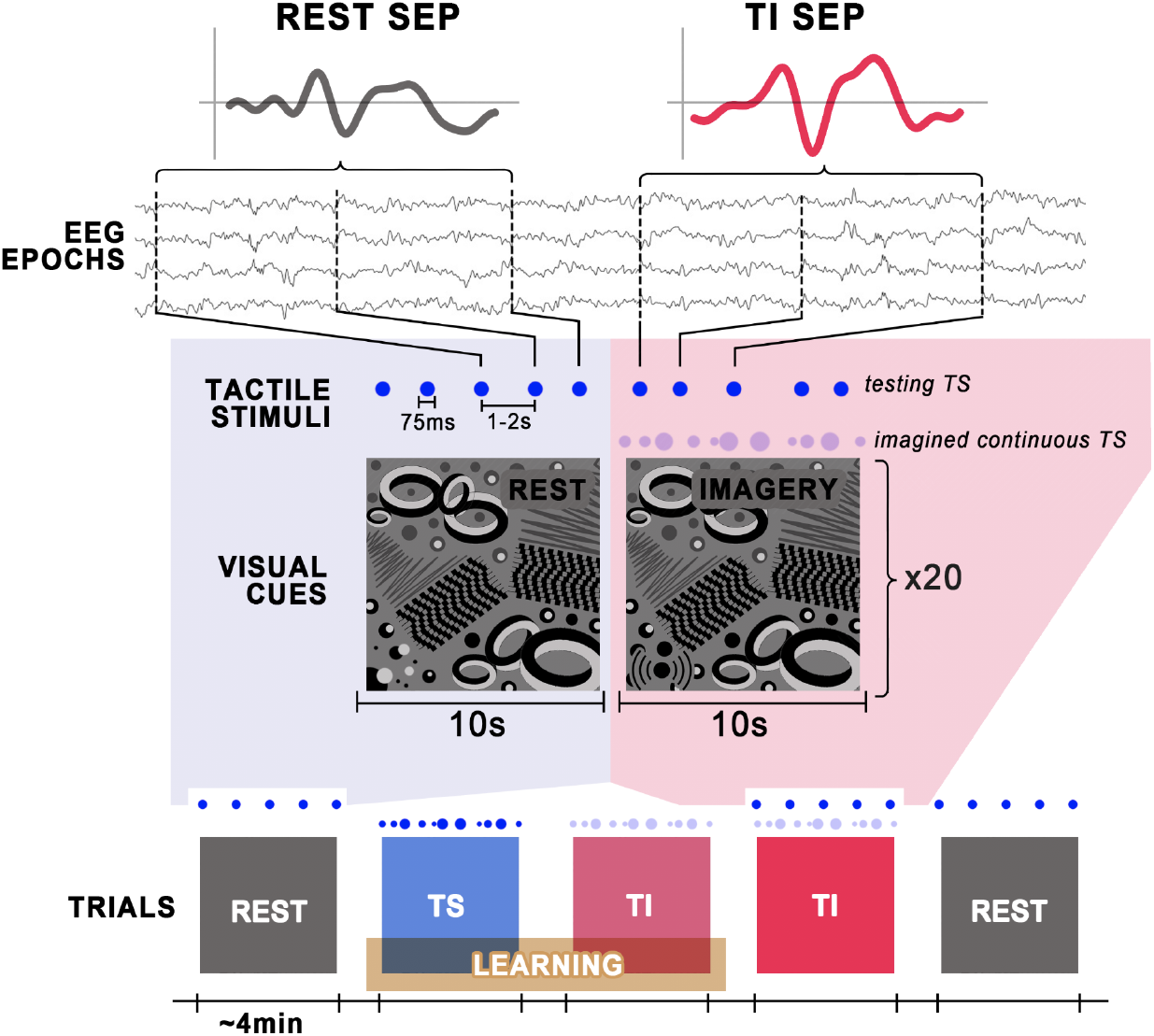
The experimental sequence. We recorded Somatosensory Evoked Potentials (SEP) in response to short (75 ms) vibrational stimuli at rest in the beginning and at the end of each experimental session (REST trials). In the experimental session, participants were familiarized with vibrostimulation of the right index fingertip (Tactile Stimulation, TS-learning trials) and then asked to mentally reproduce these sensations (Tactile Imagery, TI-learning trials). Experimental session also contained a TI condition (TI trials) with short (75 ms) vibrational stimuli. Solid blue dots represent vibrostimulation patterns. Transparent dots represent imaginary vibration patterns.

**Figure 2.**
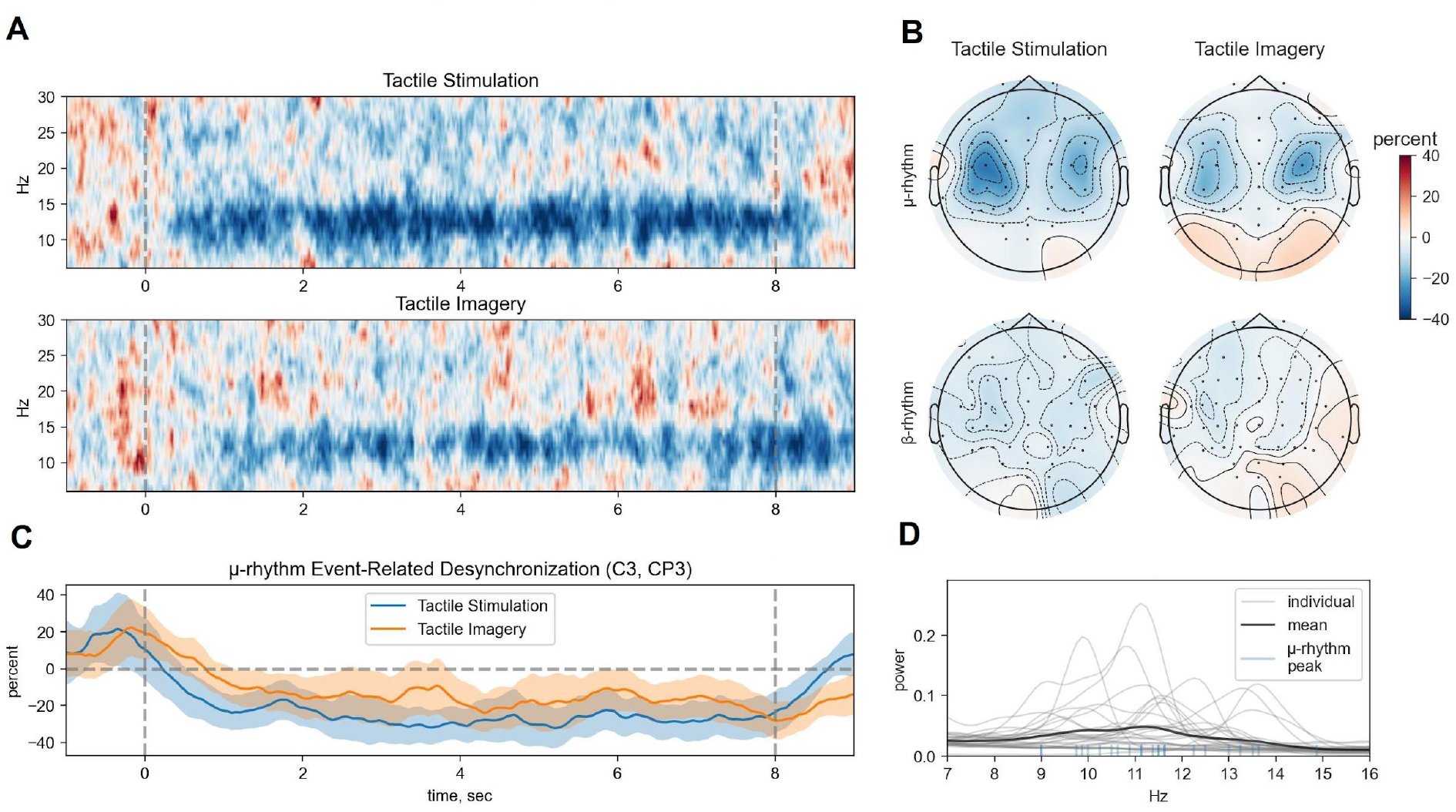
Group changes (N=29) in EEG activity during the learning session. Tactile Stimulation (TS) and Tactile Imagery (TI) trials are shown separately. **A**. Time-frequency dynamics (percent change relative to the resting state) of the EEG spectral power during TS (top) and TI (bottom). **B**. Temporal dynamics of the μ-rhythm Event-Related Desynchronization (ERD) during TS and TI. **C**. Topographic maps of μ- and β-rhythm ERD during TS and TI. **D**. Resting state spectra for the C3 and CP3 leads with marks showing individual μ-rhythm frequencies.

During TI, we also observed μ- and β-rhythm ERD with the same localization but lower intensity than during TS (*W* = 88, *p* = 0.0041). Also, despite the tendency for ERD to be stronger in the contralateral hemisphere during TI, there was no statistically significant difference between the μ-rhythm ERD in the contra- and ipsilateral hemispheres.

We observed strong positive correlation between the μ-rhythm ERD during TS and μ-rhythm ERD during TI (Spearman’s rank correlation, *r*_*S*_ = 0.731, *p* < 0.0001), that is the participants that had lower μ-rhythm ERD during TS also had lower μ-rhythm ERD during TI (Fig. 3A). We also observed correlation between the μ-rhythm power at rest and μ-rhtyhm ERD during TI (Spearman’s rank correlation, *r*_*S*_ = -0.653, *p* = 0.0001). Note that among the 29 participants, 6 people were not able to voluntarily generate detectable μ-rhythm ERD.

**Figure 3.**
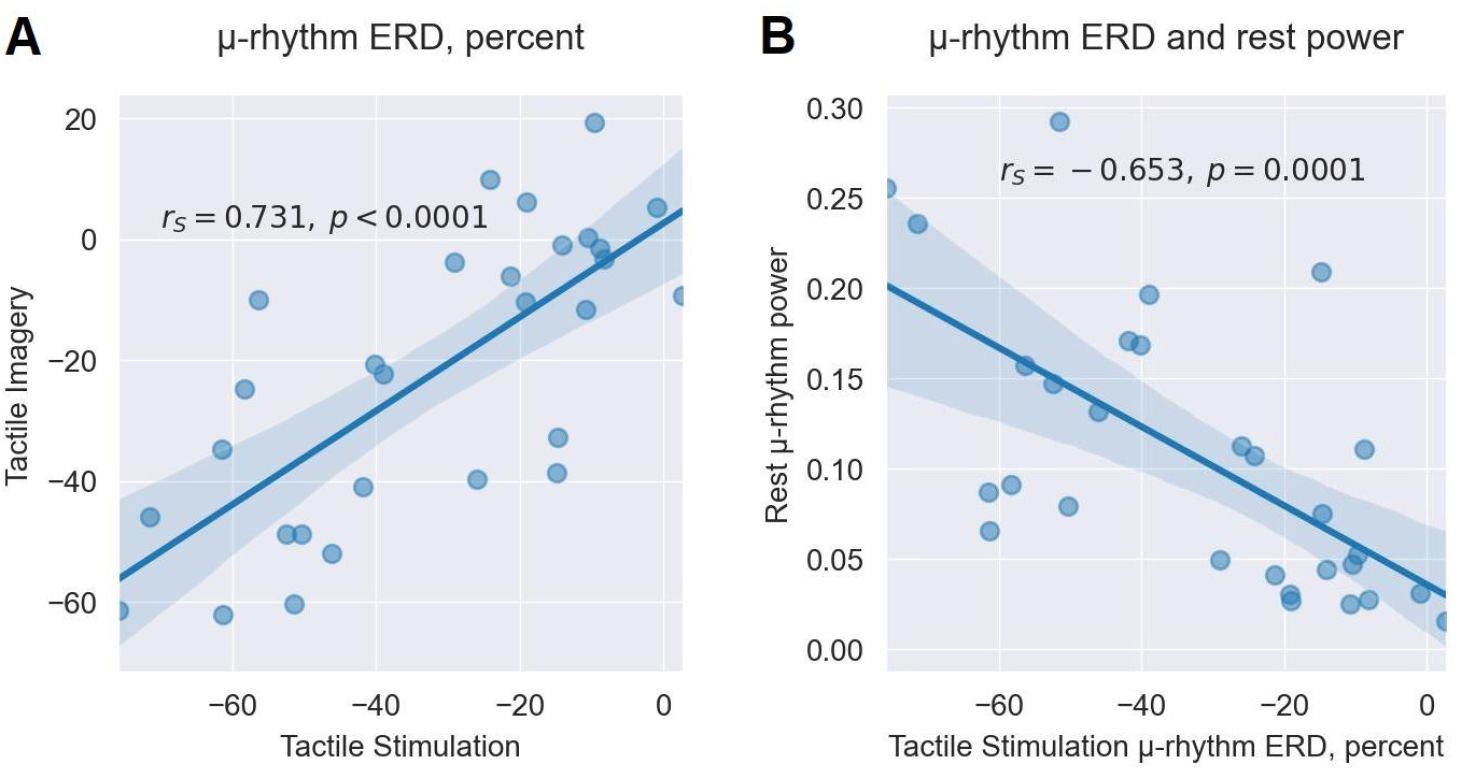
The results of the correlational analysis for (A) the μ-rhythm Even-Related Desynchronization (ERD) during Tactile Stimulation versus Tactile Imagery and (B) the μ-rhythm ERD versus its rest power.

During TS, the μ-rhythm ERD tended to start earlier but also end more abruptly than during TI (Fig. 2B). Additionally, note the difference in individual μ-rhythm frequencies across the participants (Fig. 2D).

### Event-Related Potentials (ERPs) in Response to Short Vibratory Pulses

SEPs in response to 75-ms vibratory pulses consisted of three prominent components: a P100 response in the contralateral centro-parietal region (∼102 ms latency; CP3, P1, P3, P5 leads), a P200 response in the centro-frontal region (∼229 ms latency; F3, FC1, FC3, FCz, F4, FC2, FC4 leads), and a P300 response in the contra- and ipsilateral central regions (∼301 and ∼326 ms latency in contra- and ipsilateral hemispheres, respectively; C5, CP5 and C6, CP6 leads). At rest, P100 component was also noticeable in the corresponding region of the ipsilateral hemisphere (∼133 ms latency, CP4, P2, P4, P6 leads, fig. 4, table 2), however, with statistically significantly lower amplitudes (*W*_*P100*_ = 115, *p*_*P100*_ = 0.026).

**Table 2.** Somatosensory Evoked Potential amplitudes (median [25th – 75th percentiles]).

| Component | Hemisphere | Latency, ms | Rest | Tactile Imagery |
| --- | --- | --- | --- | --- |
| P100 | Contralateral | 102 | 0.14 [0.02 – 0.20] | 0.15 [0.05 – 0.26] |
| P100 | Ipsilateral | 133 | 0.07 [–0.03 – 0.19] | 0.14 [0.09 – 0.24] |
| P200 | Frontal channels group | 229 | 0.21 [0.10 – 0.30] | 0.27 [0.18 – 0.40] |
| P300 | Contralateral | 301 | 0.11 [0.02 – 0.22] | 0.20 [0.12 – 0.27] |
| P300 | Ipsilateral | 326 | 0.07 [−0.06 – 0.19] | 0.23 [0.07 – 0.31] |

**Figure 4.**
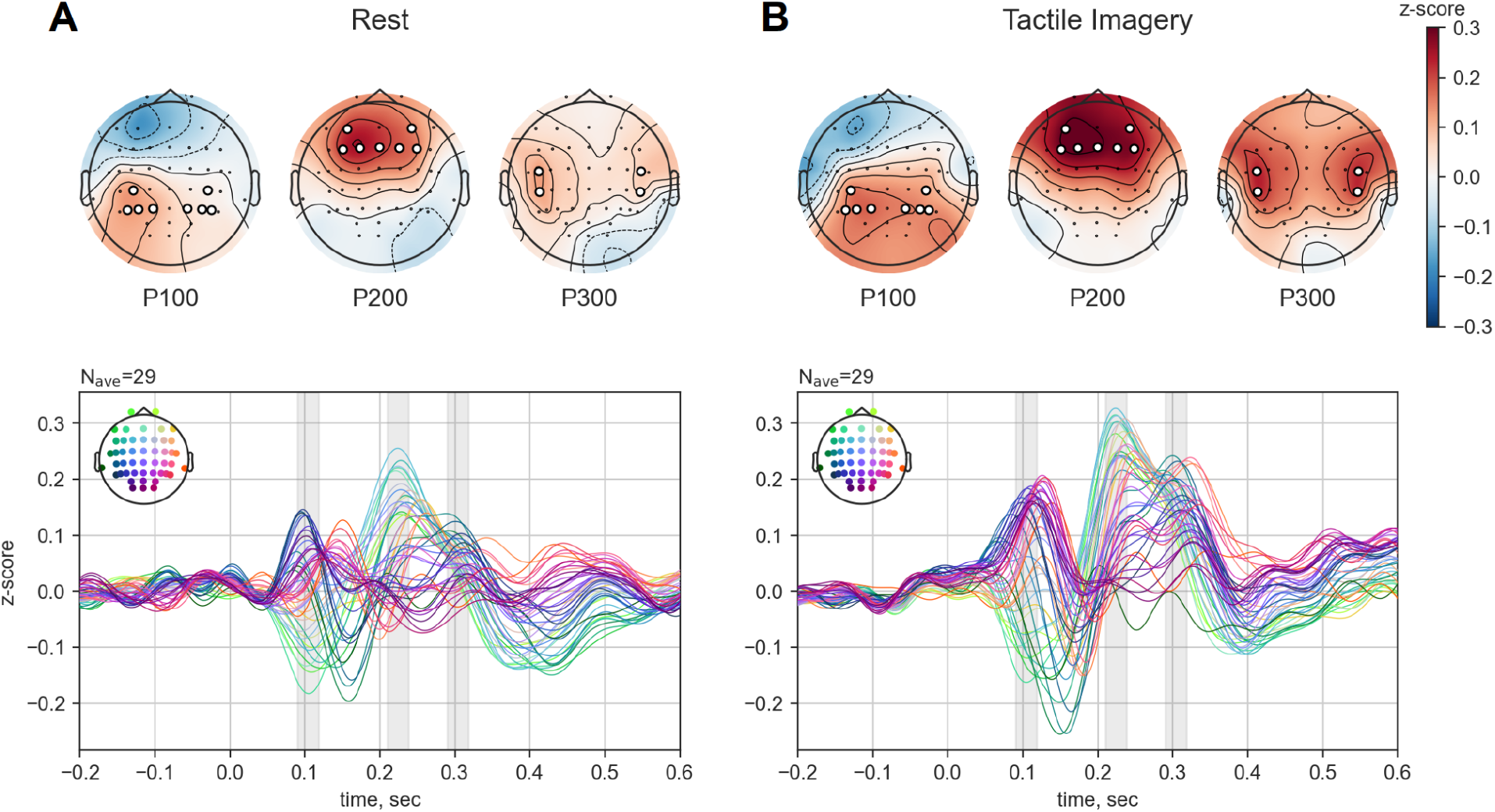
Somatosensory Evoked Potentials (SEPs) in response to short (75 ms) vibrotactile pulses and localization of their components (P100, P200, P300) at Rest and during Tactile Imagery. The grand average across subjects (N=29) is shown. The target leads for which the amplitudes of the components in the contra- and ipsi-lateral hemispheres were calculated are marked with white dots.

### Time-Frequency Responses to Short Tactile Stimuli

In the time-frequency maps for the response to 75-ms vibratory stimuli, we observed power changes in three time-frequency ranges: Event-Related Synchronization (ERS) of the θ-rhythm in contralateral centro-parietal region (3–8 Hz, 100–300 ms; Fz, F3, FCz, FC1, FC3 leads), ERD of μ- and β-rhythms (100–600 and 100-300 ms, correspondingly; C1, C3, C5, CP1, CP3, CP5 leads) in contra- and ipsilateral central regions (Fig. 5, Fig, 7A).

**Figure 5.**
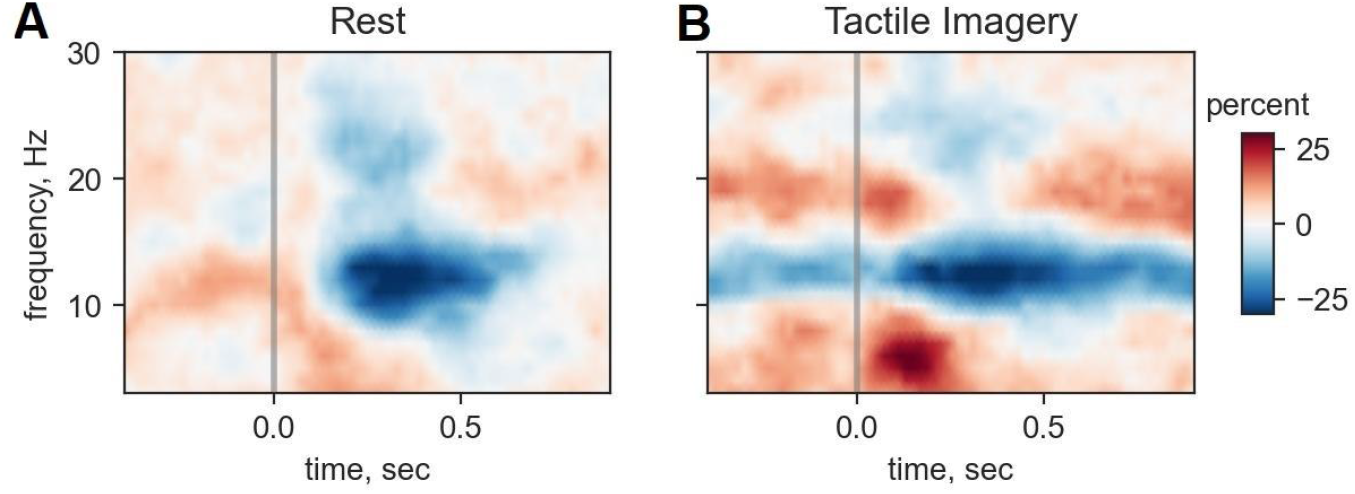
Time-frequency maps of responses to 75-ms vibratory pulses at rest (A) and during Tactile Imagery (B). Plots represent an average for the C3and CP3 leads.

### Tactile Imagery modulates ERPs and Time-Frequency responses

We recorded responses to 75-ms vibratory stimuli during three periods: TI and two rest periods (one at the beginning and the other at the end of the experimental session). We statistically analyzed the amplitudes of P100, P200, P300 peaks during these periods.

We did not find any statistically significant difference between the ERP peak amplitudes at the beginning and at the end of the experimental session. Also, there was no statistically significant difference between the ERP peak amplitudes calculated for the first and last 50 vibratory stimuli applied during the resting state. Given that there was no difference between these conditions, we combined the last 50 epochs from the resting states before and after the TI session (100 epochs in total) to obtain an average Rest ERP.

We found statistically significant increase in the P100 peak in the ipsilateral hemisphere and P200 peak in the frontal channels group during TI compared to general rest ERP (*W*_*P100*_ = 98, *p*_*P100*_ = 0.0086; *W*_*P200*_ = 112, *p*_*P200*_ = 0.021). We also found statistically a significant increase in the P300 amplitude during TI in both hemispheres compared to general rest ERP (Fig. 6; *W*_*contra*_ = 81, *p*_*contra*_ = 0.0023; *W*_*ipsi*_ = 80, *p*_*ipsi*_ = 0.0022). We also calculated LI index for each component and despite there were tendencies for LI to increase during TI for all three components, it was only statistically significant for the P200 component when it was split into two contra- (F3, FC1, FC3 leads) and ipsilateral (F4, FC2, FC4 leads) parts (*W* = 86, *p* = 0.0035).

**Figure 6.**
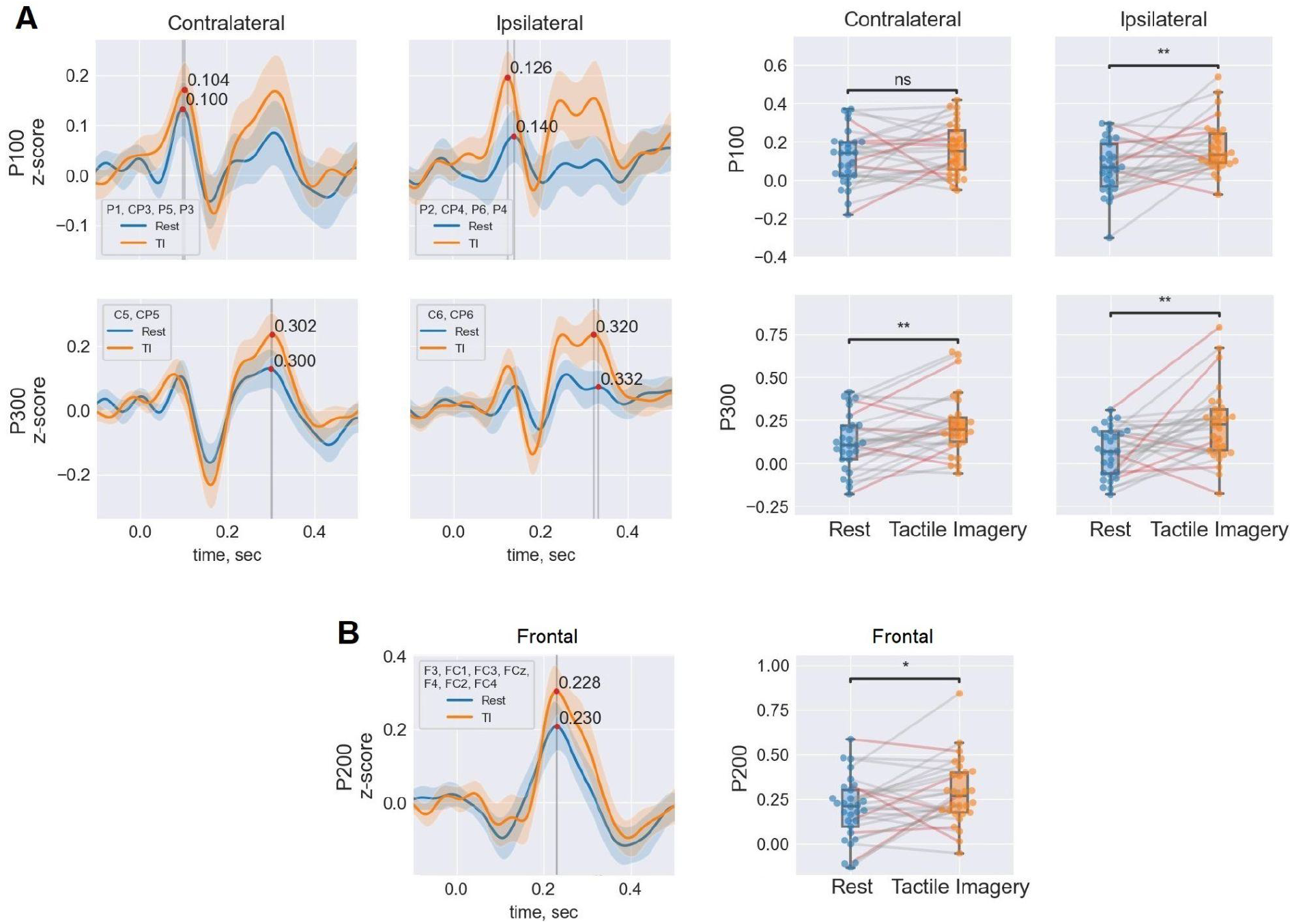
Somatosensory Event-Related Potentials to 75 ms vibrational stimuli averaged across the target leads and boxplots of the mean amplitudes for the components in contra- and ipsilateral hemispheres. In the boxplots, each point represents an individual participant, lines connect the measurements taken from the same participant, and red lines represent participants that were not able to voluntarily generate detectable μ-rhythm Event-Related Desynchronization.

We found a statistically significant increase in the θ-ERS in response to 75-ms vibratory stimuli during TI (Fig. 7, Table 3; *W* = 67, *p* = 0.0013).

**Table 3.** Powers of different spatial-temporal-frequency ranges in response to 75 ms vibratory stimuli (median [25th – 75th percentiles]).

| Time-frequency range | Rest | Tactile Imagery |
| --- | --- | --- |
| $\theta$: 100-300 ms | 0.10 [0.03 – 0.30] | 0.25 [0.14 – 0.36] |
| $\mu$: 100-600 ms | -0.16 [-0.31 – -0.02] | -0.11 [-0.29 – -0.01] |
| $\beta$: 100-300 ms | -0.08 [-0.14 – 0.03] | -0.07 [-0.16 – -0.01] |

**Figure 7.**
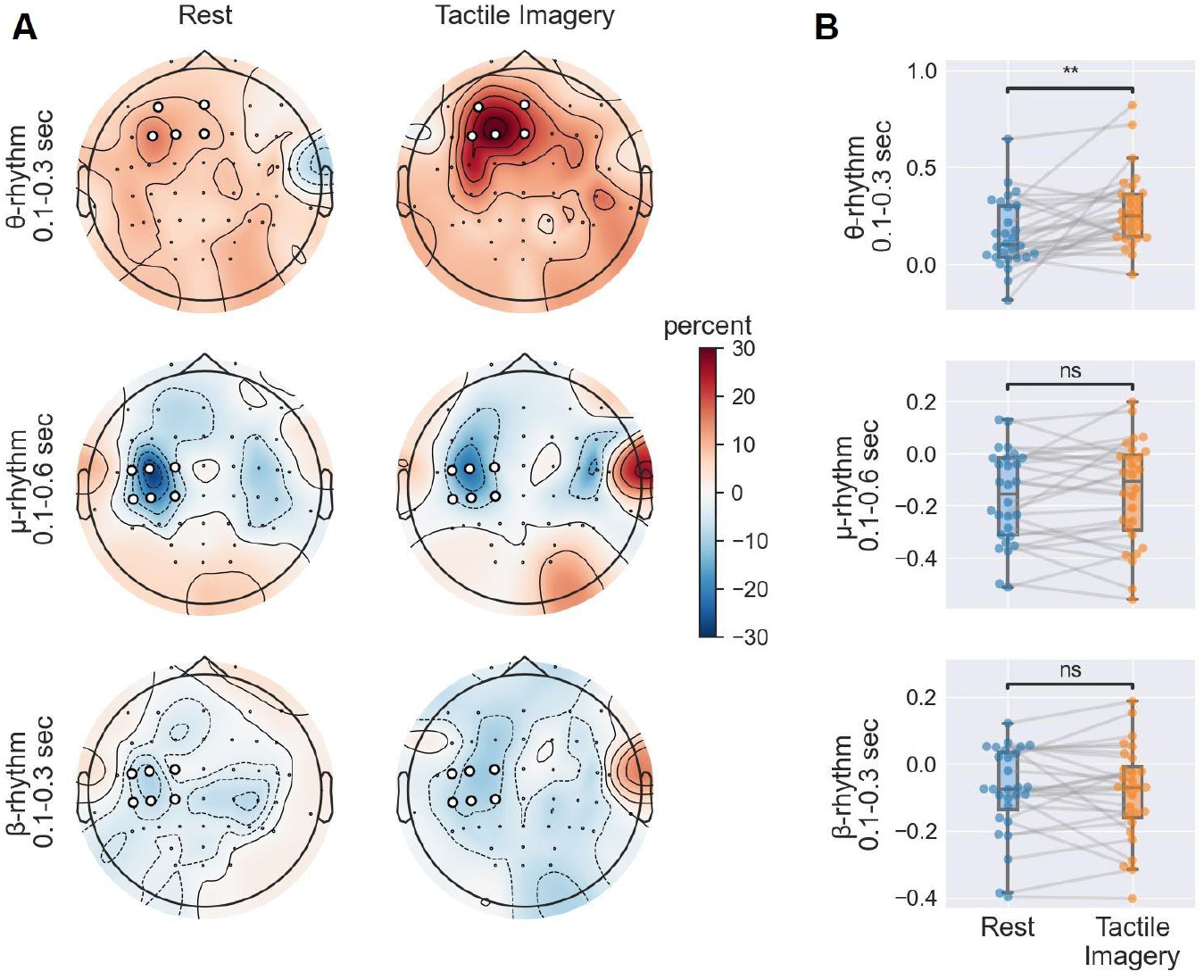
Localization of Event-Related time-frequency changes in response to 75-ms vibratory stimuli (A, target leads are marked with white dots) and boxplots of the mean power of the distinct Event-Related changes across the target leads (B).

We also found a strong correlation between P200 amplitude during TI and θ-ERS during TI (Spearman’s rank correlation, *r*_*S*_ = 0.477, *p* = 0.01), and between P200 amplitude during TI and

θ-ERS at rest (Spearman’s rank correlation, *r*_*S*_ = 0.604, *p* = 0.0007). No statistically significant correlations between P200 amplitude and θ-ERS were found for the other combinations of conditions (Spearman’s rank correlation; P200 at rest versus θ-ERS at rest: *r*_*S*_ = 0.297, *p* = 0.12; P200 at rest versus θ-ERS during TI: *r*_*S*_ = 0.126, *p* = 0.52).

## Discussion

In this study, we employed EEG recordings to examine cortical responses to tactile imagery. These EEG patterns were compared with the effects of real vibrotactile stimulation of the fingertip. Next, responses to short vibrotactile stimuli were assessed in the presence and in the absence of TI. We found that vibrostimulation and TI modulated EEG in a similar way. Moreover, TI affected the responses to short vibratory pulses. These findings clarify how imagined and real tactile sensations interact at the cortical level.

### Event related desynchronization during learning of tactile imagery

These experiments started with a session where either long (8-s) trains of vibrostimulation were applied to the fingertip or the participants imagined vibrostimulation for the same duration without actually being stimulated. During both the TS and TI, ERDs occurred in the μ-range (8-13 Hz). In the TI trials, the ERD began 0.5 seconds following the visual cue that instructed imagery start. The ERD persisted throughout the 8-s interval in both conditions (Fig. 2A, Fig. 2B). The μ-ERD was predominantly localized in the contralateral hemisphere with the strongest effects for the C3-CP3 channels (Fig. 2C). These findings are generally consistent with the previous studies where TI was found to cause desynchronization of the mu rhythm (Yao et al., 2022a; Yakovlev et al., 2023). Thus, the μ-ERD appears to be a reliable indicator of an ongoing somatosensory processing of both real and imagined tactile stimuli.

Furthermore, an across-subject positive correlation was found between the ERD values during the TI and TS conditions (see Fig. 3A), which strengthens the suggestion that imagined and real tactile stimuli are processed in a similar way and by the same cortical areas. The individual differences in the ERD values are most likely related to anatomical locations of the sensorimotor areas engaged and the specifics of the μ-rhythm in each participant. Indeed, in the MI domain, SMR-based BCI performance varies depending on individual brain structures (Halder et al., 2013; Kasahara et al., 2015). Additionally, we found that subjects with a more prominent μ-rhythm peak exhibited greater ERD, which agrees with Blankertz et al.’s (2010) suggestion made for MI that the strength of the SMR-peak during the resting state could be used as a predictor of good BCI performance during imagery. Similarly, our results suggest that ERD-based decoding could be implemented in both TS and TI-based BCIs and perhaps ERD effects could be enhanced by BCI training. Overall, our present study aligns with the findings of Yao et al. (2018, 2022b), which demonstrated that BCI performance can be enhanced using tactile-induced ERD for calibration.

### Tactile-imagery effects on SEPs

Our study revealed a significant increase in the ipsilateral P100 peak, with no changes observed in the contralateral P100. The ipsi-P100 had a delayed latency compared to the contra-P100, which suggests the involvement of transcallosal connectivity. We suggest that the P100 peak reflects the activation of the primary somatosensory cortex. This is supported by two observations: the contralateral predominance of the P100 and its relatively early timing, possibly corresponding to the P70 component reported in the studies using cutaneous electrical stimulation. A rather gradual onset of vibrotactile stimulation could explain the later peak latency compared to the response to electrical stimulation. The absence of the earlier components (P20 and P50) observed when electrical stimulation is used (Hämäläinen et al., 1990; Novičić and Savić, 2023) could be also explained by the fact that the onset of vibrotactile stimulation was not abrupt enough.

The facilitation of P100 during TI could be explained by an attention-like mechanism where TI both increases activity of the somatosensory cortex and makes it more responsive to peripheral inputs. Yet, the observed decrease in P100 latency only in the ipsilateral hemisphere is hard to explain in terms of selective attention, and it is tempting to speculate that imagining a skin area being stimulated is different from focusing attention on the same skin area and expecting a stimulus to occur. Additionally, it is important to consider the organization of S1. Specifically, the BA3a subregion of S1 is responsible for processing proprioceptive information from muscles and joints, while signals from the skin are processed in BA3b, 1, and 2 (Martuzzi et al., 2014). BA2 also represents a higher hierarchical level that integrates both skin and proprioceptive signals. Tactile stimulation was found to activate all S1 subregions in somatotopic correspondence, whereas tactile imagery mainly engages BA2 via top-down pathways (Yoo et al., 2003; Schmidt and Blankenburg, 2019). Consequently, the lack of a significant change in the contralateral P100 could be attributed to the fact that TI affects a hierarchically higher BA2 subregion.

It should be noted that the vibrotactile stimulation used in our study undoubtedly activated Pacinian afferents, which project strongly to S2 (Fisher et al., 1983). According to perceptual-grounding theory (Barsalou, 2008; Schmidt et al., 2014), the imagery of vibration-related tactile sensations should activate the corresponding S2 regions. In addition to the P100 peak, a later centrally localized positive SEP was observed in response to tactile stimulation with a latency of approximately 300 ms. The bilateral distribution and late latency of this peak suggests that it originated from S2 (Hämäläinen et al., 1990; Hoechstetter et al., 2001). The significant increase in P300 across both hemispheres during TI is consistent with the role of S2 in the perceptual grounding of TI. Interestingly, the contralateral P300 had a shorter latency than the ipsilateral, suggesting that the ipsilateral activation was not mediated by direct bottom-up inputs, but rather by transcallosal connections (Karhu and Tesche, 1999; Hoechstetter et al., 2001).

Previous studies of mental imagery (including tactile) have focused primarily on primary sensory regions, which are thought to underlie the sensory specificity of mental imagery. However, similar to our study, Yoo et al. (2003) showed that the secondary somatosensory cortex is involved in tactile imagery. In their study, TI-induced S2 activation was limited to the contralateral hemisphere. Based on this literature, we expected the current study to show a contralateral predominance of TI effects. Surprisingly, we found a greater increase in ipsilateral SEP peaks (P100 and P300). To explain the unexpected lateralization of TI effects on SEPs, we proposed several hypotheses. First, the left somatosensory areas could be overwhelmingly active during tactile stimulation, so TI did not lead to a noticeable increase in the activity of these areas. On the other hand, during TS at rest, the ipsilateral regions were little activated, and TI could have had a stronger effect and thus the amplitude of the ipsi-SEPs. Second, ipsi-SEPs are more likely to be elicited via transcallosal connections, and their peak amplitude depends on the excitability of both ipsilateral and contralateral cortical networks, which in turn could participate in interhemispheric transfer but not SEP formation. Thus, an increase in ipsi-SEPs amplitude could be elicited by the additive effects of both contralateral and ipsilateral networks. Moreover, since S2 is responsible for the intermanual transfer of tactile image while tactile learning (Ridley and Ettlinger, 1976), ipsi-S2 activation in TI may indicate the transfer of the image into the somatosensory representation of the opposite hand. This transfer may be part of the process of memorizing the tactile image that is triggered each time a tactile stimulus appears. Since working memory is involved in this process, the somatosensory P300 observed here could be related to this memorization.

Mental imagery as an attentional mechanism for reactivating tactile sensations in memory is a plausible explanation. The mental representation of stimulus features could engage attentional mechanisms involved in the selection and manipulation of sensory information during imagery (Theeuwes et al., 2011; Schmidt and Blankenburg, 2019). Studies on somatosensory attention have demonstrated activation of S2 by spatial attention to the stimulated body part (Hämäläinen et al., 1990; Karhu and Tesche, 1999). In our study, we observed an increased P300 peak during tactile imagery. However, it is difficult to claim that these effects were exclusively related to perceptually based imagery and not to attentional mechanisms. The dissociation of attentional mechanisms from neural processes reflecting the representation of mental content or “pure” mental imagery has been a topic of debate. Different process models of mental imagery (Kosslyn, 2005) argue against considering them as independent psychological constructs (Gazzaley and Nobre, 2012).

### Frontal Activation patterns modulated by Tactile Imagery and the role of Working Memory

We observed a θ-activity synchronization evoked by tactile stimuli that increased in the left frontal area as TI continued. The frontal-θ increase is known to be associated with top-down memory processes (Klimesch, 1999). Previous research suggested that the lateral prefrontal cortex (LFPC) is involved in tactile working memory (Auksztulewicz et al., 2011; Spitzer et al., 2013). Accordingly, we hypothesize that the observed θ increase in the tactile imagery condition is an indicator of activation in the lateral prefrontal cortex (LPFC), which in turn could be caused by neural processing associated with TI. This hypothesis is supported by previous studies highlighting the role of the left inferior frontal gyrus, a part of the LPFC, in TI (Schmidt et al., 2014).

Consistent with these findings, the increased θ-ERS observed in our study likely signifies activation of the cortical network involved in the processing and integrating sensory information from working memory during mental image construction. However, it is worth noting that time-locked EEG deflections could be misleadingly interpreted as event-related synchronization (ERS) in the θ-range, potentially leading to misinterpretation of θ-ERS induced by external stimuli (Harper et al., 2014). We found a frontal-localized P200 SEP peak, which could be mistakenly identified as θ-ERS on time-frequency graphs. Notably, θ-ERS showed a distinct localization in the left frontal EEG channels, whereas the frontal P200 showed a broader distribution across frontal channels in both hemispheres (Fig. 4, Fig. 7), and correlation analysis did not reveal any significant associations between θ-ERS and P200 peak amplitude at rest (Spearman’s rank correlation; P200 at rest versus θ-ERS at rest: r_S_ = 0.297, p = 0.12).

We observed a statistically significant increase in the P200 peak during the tactile imagery (Fig. 4), which also could be related to LPFC activation during TI. Literature on mental imagery suggests LPFC to be involved in imagery regardless of the content being imagined (Ishai et al., 2000; Yoo et al., 2001; McNorgan, 2012; Schmidt et al., 2014; Pearson, 2019). It was proposed that LPFC is involved in the active construction of a mental image. Schmidt et al. (2014) suggested that imagery could be considered as a dynamical component of working memory. Based on our findings and the reviewed literature, we propose that TI facilitates the processing of tactile information within the prefrontal cortex, which is manifested by the increased θ-ERS and P200 (Fig. 8).

**Figure 8.**
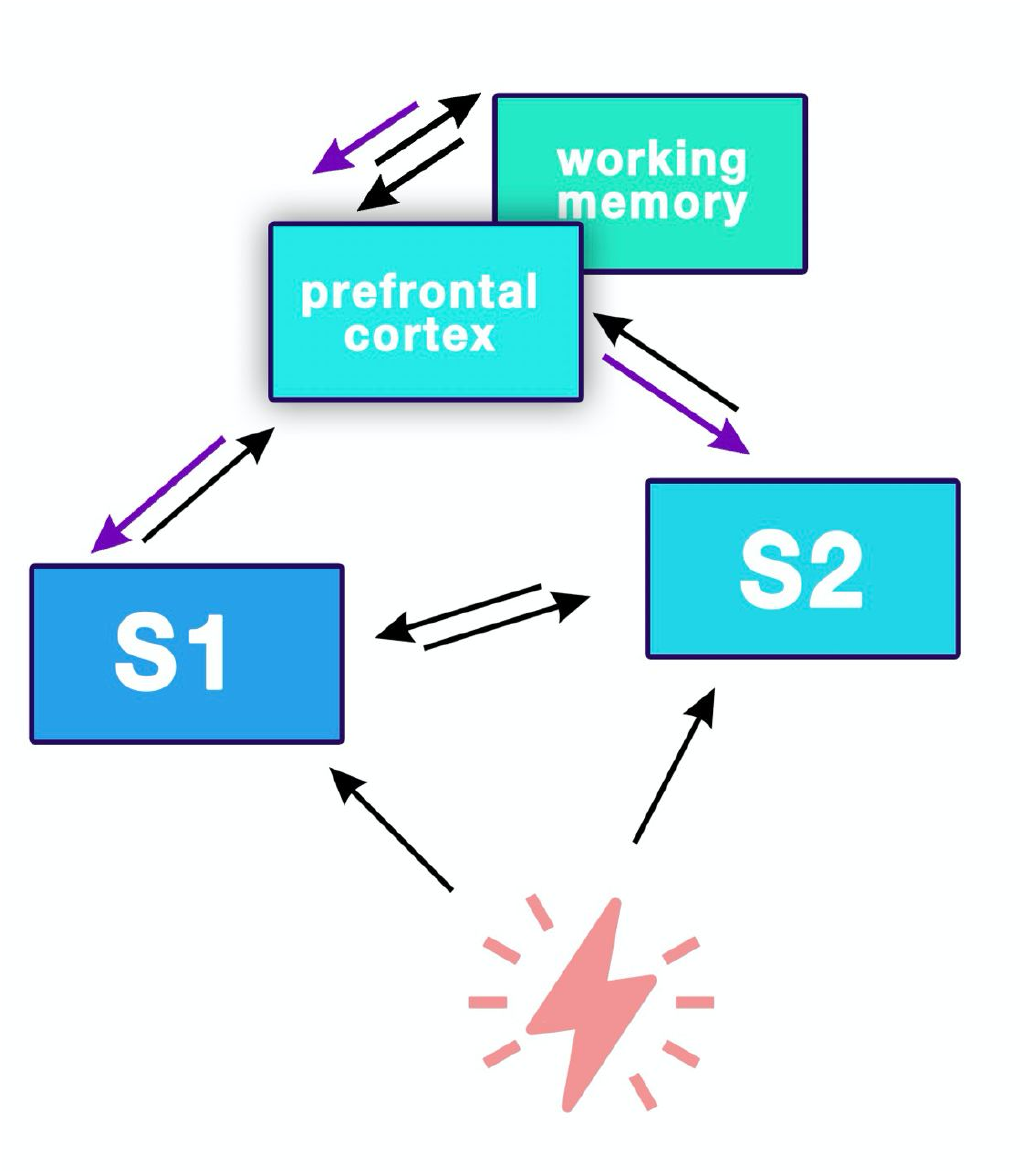
Schematic representation of the imagery process that we propose based on the obtained results. Black arrows represent normal somatosensory processing. Blue arrows represent facilitation of the processing during tactile imagery.

## Acknowledgments

This work is supported by the Russian Science Foundation under grant №21-75-30024.

## Notes

**Conflict of interest statement** Authors report no conflict of interest.

https://github.com/MarkaMorozova/Tactile-Imagery

### Competing Interest Statement

The authors have declared no competing interest.

